# Nanoscale clustering by O-antigen-Secretory Immunoglobulin-A binding limits outer membrane diffusion by encaging individual *Salmonella* cells

**DOI:** 10.1101/2023.07.13.548943

**Authors:** Alyson Hockenberry, Milad Radiom, Markus Arnoldini, Yagmur Turgay, Matthew Dunne, Jozef Adamcik, Beth Stadtmueller, Raffaele Mezzenga, Martin Ackermann, Emma Slack

**Affiliations:** Department of Environmental Systems Science, D-USYS, ETH Zurich, Zurich, Switzerland; Department of Environmental Microbiology, Eawag, Dubendorf, Switzerland; Department of Health Sciences and Technology, D-HEST, ETH Zurich, Zurich, Switzerland; Department of Biochemistry, Center for Biophysics and Quantitative Biology, and Carl R. Woese Institute for Genomic Biology, University of Illinois, USA; EPFL, School of Architecture, Civil and Environmental Engineering, Lausanne, Switzerland; Botnar Research Center for Child Health, Basel, Switzerland

## Abstract

Secreted immunoglobulins, predominantly SIgA, influence the colonization and pathogenicity of mucosal bacteria. While part of this effect can be explained by SIgA-mediated bacterial aggregation, we have an incomplete picture of how SIgA binding influences cells independently of aggregation. Here we show that akin to microscale crosslinking of cells, SIgA targeting the *Salmonella* Typhimurium O-antigen extensively crosslinks the O-antigens on the surface of individual bacterial cells at the nanoscale. This crosslinking results in an essentially immobilized bacterial outer membrane. Membrane immobilization, combined with Bam-complex mediated outer membrane protein insertion results in biased inheritance of IgA-bound O-antigen, concentrating SIgA-bound O-antigen at the oldest poles during cell growth. By combining empirical measurements and simulations, we show that this SIgA-driven biased inheritance increases the rate at which phase-varied daughter cells become IgA-free: a process that can accelerate IgA escape via phase-variation of O-antigen structure. Our results show that O-antigen-crosslinking by SIgA impacts workings of the bacterial outer membrane, helping to mechanistically explain how SIgA may exert aggregation-independent effects on individual microbes colonizing the mucosae.

## Introduction

Mucosal surfaces are rich in the secretory immunoglobulins, predominantly SIgA, which protect host tissues from would-be invasive microbes. In the case of non-Typhoidal *Salmonella*, recent work indicates the importance of SIgA-mediated bacteria aggregation, via both enchained growth and classical agglutination, in inhibiting bacterial interaction with the epithelium^1,2^. However, previous studies with O-antigen-targeting monoclonal antibodies have indicated the potential for SIgA to directly block bacterial motility, even in the absence of aggregation, and to uncouple bacterial membrane potential, via unknown mechanisms^3–5^. Additionally, antibody binding to bacterial surface antigens influences bacterial transcription^6,7^. A number of experimental models now support the idea that SIgA shapes microbiota composition, and is capable of both reducing and enhancing colonization levels of particular species^8–11^. Thus, while cellular aggregation is now well-understood, a range of IgA-driven phenotypes cannot be explained by our current understanding of how IgA interacts with the surface of individual bacterial cells.

In the case of *Salmonella enterica* subsepcies *enterica* serovar Typhimurium, oral-vaccine-induced protective SIgA targets the O-antigen of lipopolysaccharide^1,12^. This repetitive glycan polymer densely coats the bacterial outer membrane, largely masking non-protruding bacterial surface proteins (e.g. outer membrane porins). IgA-mediated protection against *S*. Typhimurium *in vivo* is O-antigen dependent and correspondingly O-antigen-specific monoclonal antibodies mimic the effects of a whole-cell vaccine-induced polyclonal SIgA response.

Due to its dimeric structure, each SIgA has four antigen binding domains (Fabs) and it is this high-avidity property that allows it to link between bacterial cells, generating aggregates that are resilient to high turbulent forces^1,2^. We reasoned a similar phenomenon might occur at the nanoscale: when all four Fab domains bind to an individual cell, SIgA would link O-antigens into clusters. As the O-antigen is arrayed around abundant outer-membrane porins^13^, this has the potential to “lock” O-antigens around these protein structures, generating a cage, and ultimately altering bacterial outer membrane properties.

Here we use a recombinant secretory IgA, STA121, that binds with high affinity to the O:12-0 epitope of the *S*.Typhimurium O-antigen to investigate the influence of SIgA binding on bacterial outer membrane biology. This demonstrated an essentially complete loss of O-antigen diffusion in the outer membrane upon IgA binding. This not only has consequences for membrane-fluidity-dependent cell processes but also contributes to the efficiency of aggregation. IgA-encaged cells display a polar bias of IgA-bound O-antigen, which is consistent with “locking” of lipopolysaccharide around non-diffusable OMPs, and BAM-dependent insertion of new OMPs at the cell middle. While this could enhance aggregation in a monophasic population in the presence of abundant SIgA, the same process is expected to accelerate escape from IgA-mediated aggregation, as the new phase-varied O-antigen-structure will accumulate at the point of septation, while the old IgA-bound O-antigen remains concentrated at the oldest poles.

## Results and discussion

### SIgA binding the O-antigen essentially immobilizes the outer membrane structures

In order to examine how O-antigen-binding SIgA distributes on the cell membrane, we carried out atomic force microscopy of *S*.Typhimurium coated, or not, with STA121 recombinant SIgA. To enable high-resolution imaging, the prepared *S*. Typhimurium were gently dried onto the imaging surface and imaged in ambient air, which results in a corrugated appearance of the normally “carpet-like” O-antigen (Fig 1A). When bacterial were pre-incubated with STA121 SIgA prior to imaging, the globular domains of bound SIgA were visible surrounding clumped O-antigen. (Fig 1B).

**Figure 1.**
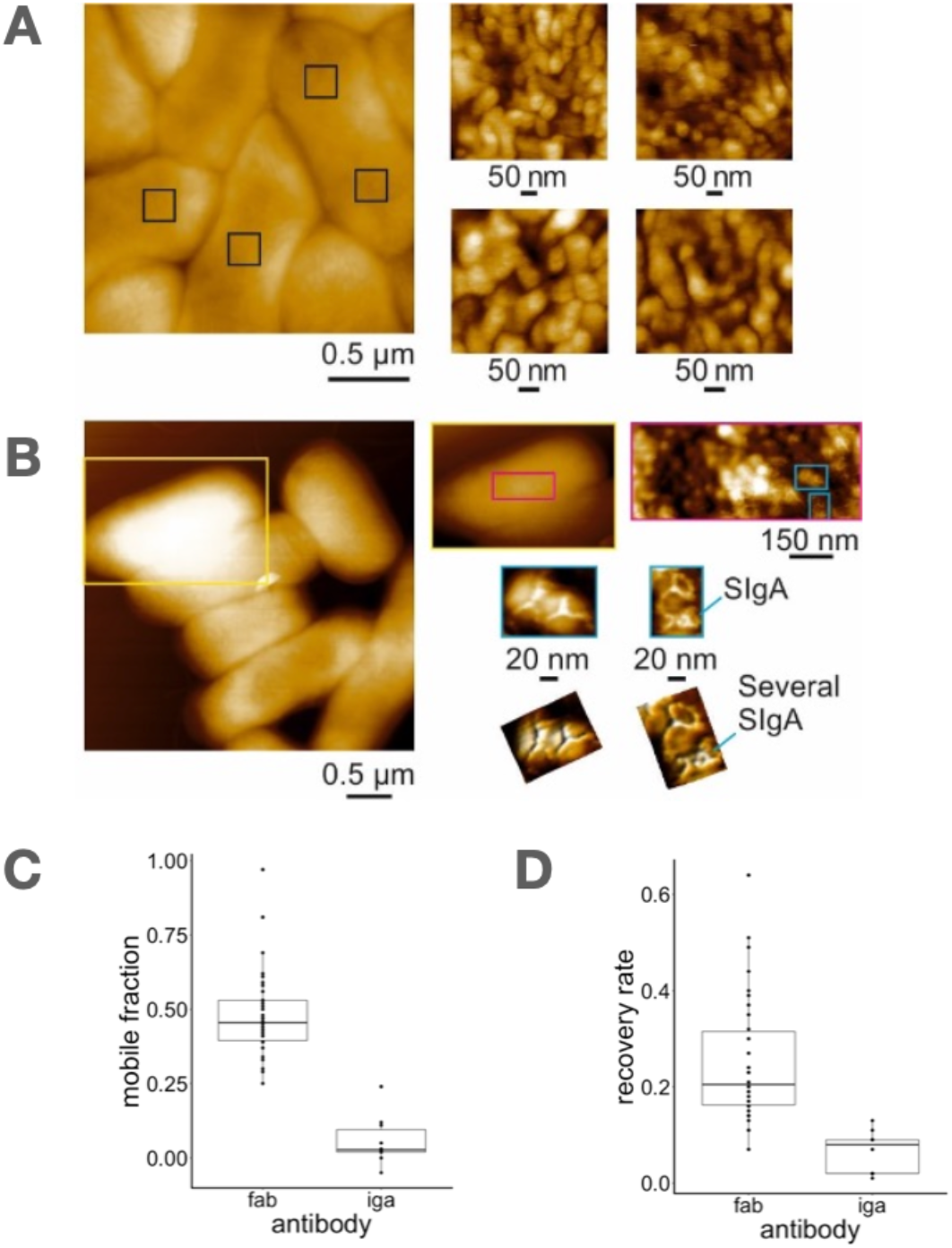
SIgA impacts the fluidity of the bacterial outer membrane. High resolution AFM height images of **A** *Salmonella* Typhimurium and **B** *Salmonella* Typhimurium exposed to SIgA. Enlarged areas are marked with boxes. Localized SIgA on O-antigen in are marked. **C** Mobile fraction and **D** recovery rate of Fab:O-antigen or SIgA:O-antigen complexes during fluorescence recovery after photobleaching experiments.

Because SIgA appeared to clump the O-antigen on the cell surface, we next measured the impact of SIgA on lateral mobility of O-antigen on the bacterial surface using fluorescence recovery after photobleaching (FRAP). In order to investigate the importance of O-antigen crosslinking in altered diffusion properties, we also generated recombinant FAb fragments of the same antibody, which binds but cannot crosslink.

The antibodies were fluorescently labeled, coated onto the surface of live *S*.Typhimurium and then a fraction of the cell surface was photobleached. Recovery of fluorescence was monitored over 5-15 minutes (Fig. 1C and D). FAb-O-antigen complexes largely retained motility, recovering with the expected kinetics. Note that due to the small size of bacteria, a major fraction of surface flurophores are bleached during this process and we do not expect 100% recovery in unmanipulated cells. In contrast, there was almost no recovery of fluorescence after photobleaching of the SIgA-O-antigen complexes, and the fraction that recovered was very slow, indicating that SIgA-O-antigen complexes are fixed in position in the outer membrane (Fig. 1C, 1D, Fig S1).

### SIgA drives biased inheritance of O-antigen through its high avidity and interaction with OMPs

Fixation of SIgA-O-antigen complexes on the outer membrane suggested that SIgA-binding may have consequences for the distribution of these complexes during bacterial growth. Of note, *Salmonella* colonization can evade adaptive immune responses via structural variation of its O-antigen through phase variation or mutation of biosynthetic genes^14,15^. A change in O-antigen structure results in loss of antibody reactivity and ultimately evasion of host immunity^12^.

However, altering a biosynthetic pathway for an abundant surface glycan does not instantaneously change the surface antigenicity. Rather, a *Salmonella* cell inserts new O-antigen into its outer membrane as it grows, resulting in gradual out-dilution of the “old” O-antigen type. Upon O-antigen structural variation, the cell will contain different O-antigen structures. Because O-antigen is highly diffusible^16^ and it is thought to be inserted uniformly across the cell envelope^17^, it is expected that the previous O-antigen structure would dilute out uniformly through growth. In order to examine the consequences of constrained O-antigen-SIgA-complex motility (Fig 1B, C), on the distribution of these complexes during bacterial growth, we fully coated *S*.Typhimurium with fluorescent STA121 SIgA and observed the pattern of fluorescence during growth using time-lapse microscopy in microfluidic devices.

Contrary to the null hypothesis of uniform mixing on the membrane and therefore uniform dilution of SIgA:O-antigen complexes, we observed enrichment of these complexes at the poles of the cells (Figs 2A). Over several generations, the SIgA-O-antigen complexes were maintained at the oldest pole cells (Fig 2A,B,C,S2, Video 1). On average, old-pole cells inherited roughly 60% of the IgA:O-antigen complexes. This biased inheritance of IgA:O-antigen complexes relied upon the multivalent nature of SIgA, as cells grown in the presence of the Fab fragment showed uniform distribution of Fab:O-antigen complexes across daughter cells, in line with uniform mixing in the outer membrane (Fig 2B,C). An identical pattern could be observed when STA121 SIgA was continuously present during culture but we used an *S*.Typhimurium strain capable of switching to the non-recognised O:12-2 O-antigen structure by DAM-dependent phase variation (refs).

**Figure 2.**
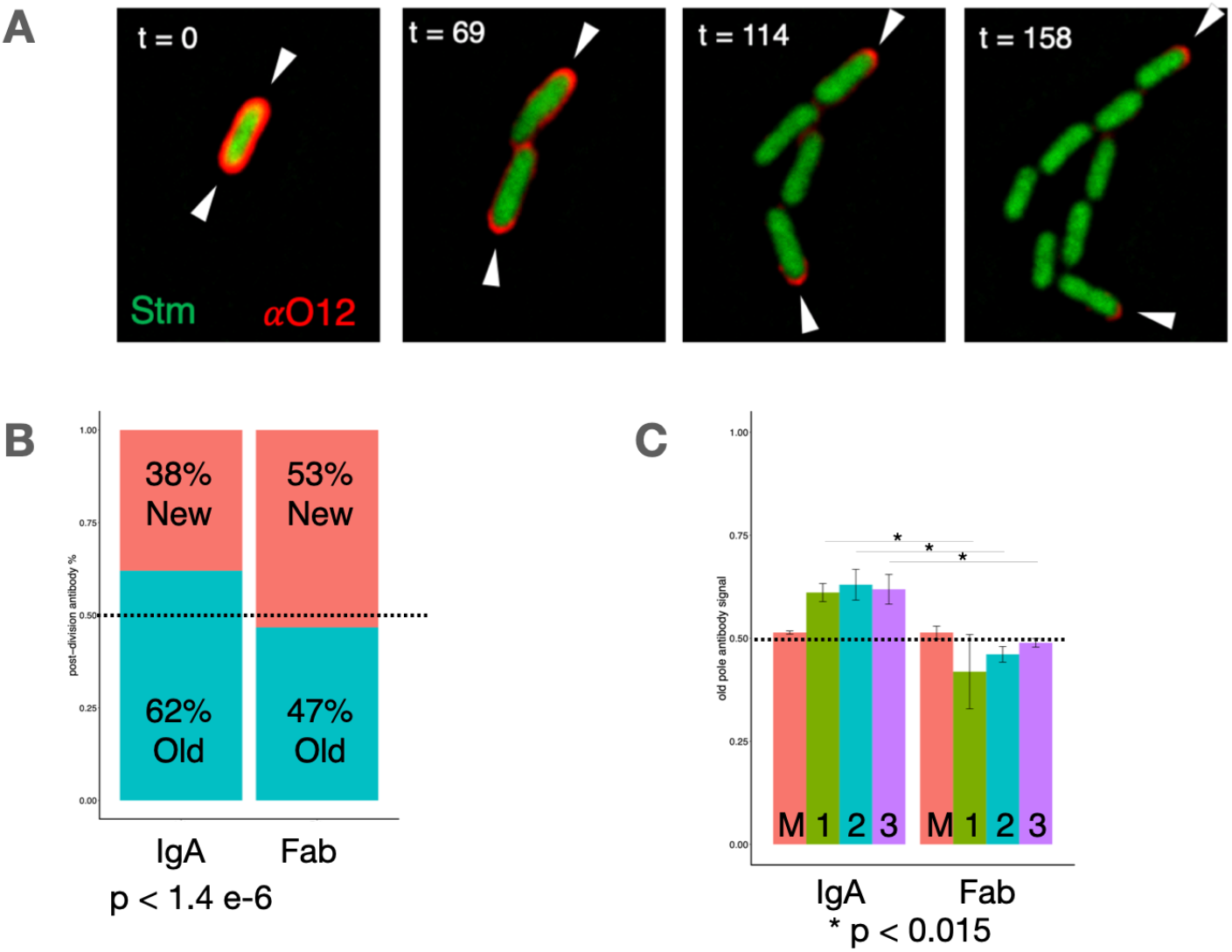
High avidity SIgA imposes biased inheritances of O-antigen. **A** Time-lapse microscopy of Salmonella cell (green) growing in the presence of fluorescently labeled SIgA (red) and undergoing O-antigen structural variation. White arrow heads indicate the oldest poles of the lineage. **B** The partitioning constant of SIgA:O-antigen or Fab:O-antigen complexes. This was quantified as fluorescence inherited by the old-pole divided by the total cellular fluorescence. SIgA vs Fab, p < 1.4 e-6 **C** Amount of SIgA:O-antigen or Fab:O-antigen inherited by daughter cells over generations. M, mother cell, 1, 1st division, 2, 2nd division, 3, 3rd division. SIgA vs Fab, *, p < 0.015 for generations 1-3

We next examined the mechanism driving polar accumulation of SIgA-O-antigen complexes. An initial hypothesis was based on the increased mass of SIgA-O-antigen compared to free O-antigen, which may be predicted to particularly strongly slow the diffusion rate on the curved surfaces of the poles. We therefore quantified the diffusion rates of SIgA:O-antigen complexes at curved and straight regions of the cells and correlated these values to local curvature (Fig S3). These analyses indicated local curvature does not correlate with the diffusion rate of SIgA:O-antigen complexes, and we could therefore reject this hypothesis.

As O-antigen is added all over the cells, the biased inheritance of O-antigen-SIgA complexes is not well explained by O-antigen addition alone. Intriguingly, other outer membrane components are reported to exhibit biased inheritance^18,19^. Older outer membrane proteins (OMPs) accumulate at the poles through a combination of two mechanisms: first, OMPs are preferentially inserted at the center of the cell^20^; and second, promiscuous protein interactions between OMPs results in gradual accumulation into large protein islands with decreased diffusivity^18^. A recent report shows OMPs are surrounded by a layer of LPS, implying OMPs and O-antigens are in very close apposition in the membrane^13^. We speculated that large SIgA:O-antigen complexes behave similarly to OMP protein islands, causing SIgA:O-antigen complexes to “hitchhike” toward the poles. Consistent with this idea, the partitioning constant of SIgA:O-antigen is similar to those reported for the outer membrane porins BtuB and TolC in *E. coli* ^18,19^(Fig 2B,C).

We first tested whether IgA:O-antigen complexes localize near OMPs. Sequential staining of *Salmonella* cells with SIgA followed by a fluorescently labeled phage tail fiber specific for OmpC indicated SIgA significantly reduces OmpC accessibility, consistent with close physical association (Fig 3A). Intriguingly, SIgA, did not limit accessibility to an LPS-core-binding phage tail fiber, suggesting either more exposure of this epitope in SIgA-bound cells. To further test the hypothesis the OMP-association was driving biased inheritance, we quantified the IgA:O-antigen partitioning constant in *Salmonella* cells lacking one component of BAM complex machinery, BamB. Cells deleted of *bamB* remain capable of OMP production and insertion, but at a much lower level than wildtype cells^20^ (Fig S4). In line with our hypothesis, cells deleted of *bamB* did not exhibit biased partitioning of SIgA-bound O-antigen, with the distribution looking similar to Fab-fragment-stained cells (Fig 3C, S5).

**Figure 3.**
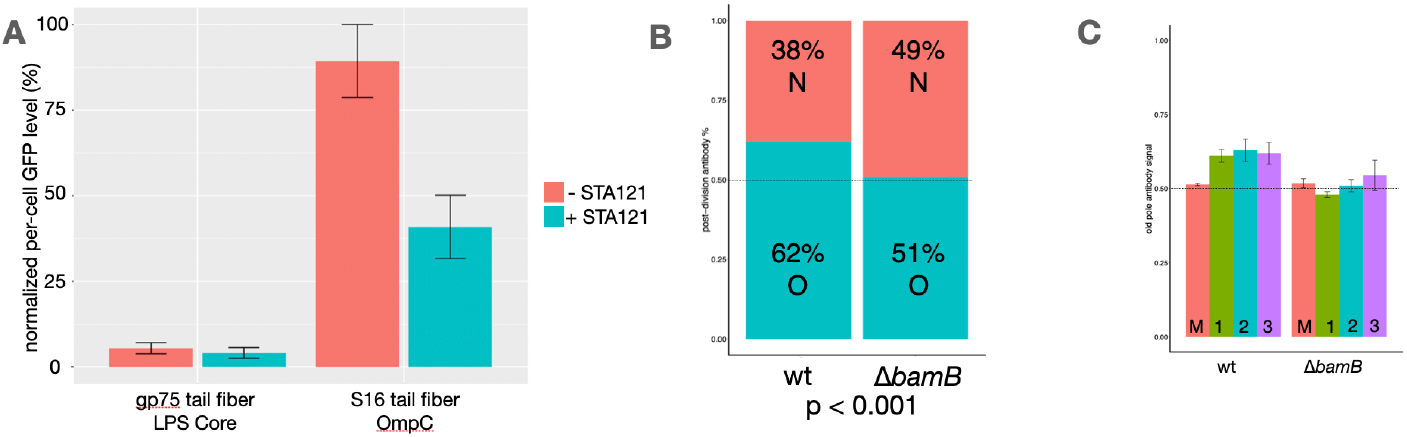
Biased inheritance of OMPs drives biased inheritance of SIgA:O-antigen. **A** Cells were incubated in the presence or absence of SIgA, washed, and then co-incubated with a fluorescently labeled marker for OmpC (S16) or the LPS core (gp75). Cells were examined by fluroescence microscopy to quantify the accessiblity of OmpC or LPS core in the presence or absence of SIgA. **B** The partitioning constant of SIgA:O-antigen in wild type or bamB cells. This was quantified as fluorescence inherited by the old-pole divided by the total cellular fluorescence. SIgA vs Fab, p < 0.001 **C** Amount of SIgA:O-antigen inherited by wild-type or bamB daughter cells by generation. M, mother cell, 1, 1st division, 2, 2nd division, 3, 3rd division.

Biased partitioning of IgA:O-antigen complexes, therefore, is driven by two factors: high-avidity binding of surface antigen and the interactions between IgA:O-antigen complexes and OMPs. Our data support a model in which diffusion limited IgA:O-antigen complexes accumulate at the oldest poles through promiscuous protein interactions with OMPs^13,18–20^.

### Biased inheritance enhances the rate of immune escape

These observations generate a model where SIgA binding to O-antigens generates a “cage” around the bacterial cells that prevents a large fraction of outer membrane molecules from freely diffusing on the bacterial surface. This is expected to be sufficient to prevent chemotaxis of IgA-bound cells, and at high levels to result in inefficient function of some outer membrane porins, which may explain previously-observed loss of proton-motive force. However, we also hypothesized that biased inheritance of O-antigen-IgA complexes may influence bacterial aggregation, and escape of this aggregation by phase-variation.

Under a model of unhindered membrane fluidity, daughters of *Salmonella* cells undergoing O-antigen structural variation would equally inherit the “old” immunogenic O-antigen and the unrecognized “new” modified O-antigen. Due to SIgA imposed biased inheritance, cells recognized by the antibody yield daughters of two types: old pole cells enriched in the immunogenic O-antigen and new pole cells enriched in the unrecognized O-antigen. Because SIgA physically accelerates the emergence of daughter cells with different antigenicity, SIgA-imposed biased inheritance enhances the rate at which fully SIgA-free cells emerge.

Because the wild-type and *bamB* mutant have different partitioning constants, we directly tested the impact of biased inheritance in an *in vitro* model of immune escape. This is based on the observation that SIgA protects the gut from *S*.

Typhimurium infection primarily through aggregation. As an individual cell within an aggregate loses antibody reactivity due to O-antigen structural variation, the strength of aggregation should weaken as the number of cross-links between cells decreases. Biased inheritance should further weaken the aggregate, by introducing daughter cells with even lower amounts of immunogenic O-antigen.

To test this, we compared aggregate sizes between the two *S*. Typhimurium strains that were IgA-coated, washed and then allowed to grow for ∼ 5 divisions, mimicking IgA-out-dilution due to phase-variation. Aggregate strength was tested by quantifying aggregate size under increasing levels of shear force (Fig 4A,B). In the absence of force, wt and *bamB* cells showed the same distribution of aggregate sizes. However, under all levels of shear force, wild-type cells showed faster disaggregation compared to the *bamB* mutant. This is consistent with the idea that biased inheritance enhances the rate of aggregate breakage, and by implication immune escape on phase-variation.

**Figure 4.**
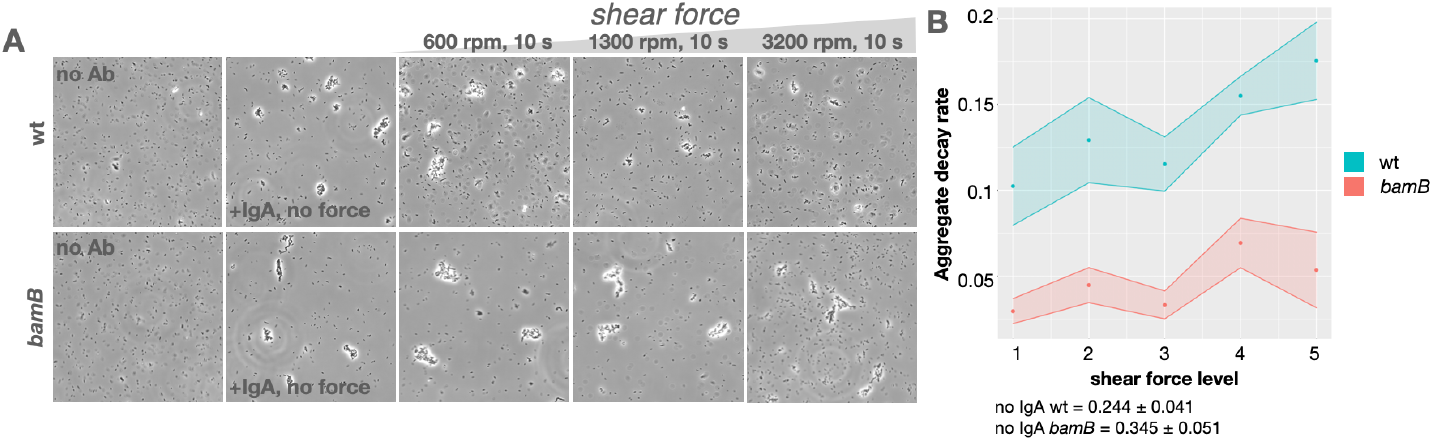
SIgA imposed biased inheritance enhances immune escape. **A** Images of wild-type and *bamB* cells grown in the presence or absence of SIgA and exposed to increasing levels of shear force. 1, 600 rpm; 2, 900; 3, 1300 rpm; 4, 2300 rpm; 5 3200 rpm. **B** Aggregate decay rate, k, as a function of shear force by strain.

Taken together, SIgA imposed biased inheritance hastens the emergence of cells devoid of immunoreactive O-antigen, enhancing the rate of immune escape by individual cells. In populations of cells, SIgA imposed biased inheritance limits the protective power of SIgA by decreasing its ability to efficiently aggregate cells undergoing O-antigen structural variation, favoring bacterial persistence in the face of an effective adaptive immune response.

### Final remarks

Our study demonstrates that SIgA binding can impact individual cells by modulating outer membrane properties. These effects on individual cells scale to effects on the population by enhancing the rate of phase variable antigen dilution during growth, and by implication accelerating immune evasion.

Several groups have reported that antibody binding decreases swimming motility and skews bacterial gene expression^5,21,22^. Our observations on the impact of SIgA on outer membrane properties provide a putative mechanism underlying these observations as relocation of flagella and chemotaxis receptors clearly requires protein movement in the outer membrane and OMP function may be broadly impacted either by direct impedance by the complexes or by mis-localization.

IgA directly impacts microbiota composition, and depending on the model used and the host, large fractions of the microbiota appear to be IgA coated in steady-state^9,23^. Though the mechanisms remain largely unclear, IgA can promote both inclusion and exclusion of species in the gut microbiota. In addition to physical aggregation mechanisms at the micrometer scale, we propose that IgA can act on microbiota composition by impacting the fitness of recognized cells at the nanometer scale. We observe that O-antigen-binding by dimeric SIgA strongly affects outer membrane properties. This change may ultimately alter bacterial growth and survival, potentially through modulation of nutrient uptake or metabolite efflux^21,24^, leading to changes in relative abundance of a given species. While we are certain that much remains to be discovered, nano-scale surface antigen aggregation and bacterial encaging may help to explain the seemingly contradictory effects of IgA on microbial colonization of mucosal surfaces.

It is also critical to point out the limitation of this study. Our study investigated the impact of a single IgA specificity, isolated from a chronically-infected human patient, with high-affinity for the O:12-0 epitope of the *S*.Typhimurium O-antigen. In this context, we have not been able to investigate the influence of high versus low affinity antibodies, nor to investigate antibodies with specificities for different parts of the *S*.Typhimurium O-antigen. Further studies should investigate how different IgA affinities and specificities affect outer-membrane fluidity, also in more diverse Gram-negative bacteria. An interesting case would be to investigate whether IgA targeting capsular polysaccharides, that are only weakly anchored in the outer membrane via diacyl lipids, can also drive this phenomenon. It would also be intriguing to investigate whether differences in antibody affinity and/or binding modality altered the transcriptional response of bacteria to IgA-targeting.

Antibody-based therapeutics are coming into view as a means of microbiota precision editing. Engineered IgA responses that can directly target bacterial properties, to ultimately enhance or inhibit their colonization, can be a powerful tool to tailor our microbiotas toward a healthful state. Understanding their mechanisms of action is essential to design robust treatments with reliable beneficial outcomes.

## Methods

### Bacterial cultivation, genetic manipulation, and recombinant IgA

Throughout the study, we used derivatives of *Salmonella enterica* subsp. enterica serovar Typhimurium, strain SB300 (a spontaneous Streptomycin-resistant SL1344 derivative). Cryostocked strains were streaked on to Lysogeny Broth (LB) Miller agar for single colonies. After 24 h, a single colony was transferred to 5 mL fresh LB in a 15 mL round-bottom tube and incubated at 37ºC with shaking at 200 rpm. A 16-18 h overnight culture prepared in this way was the starting point for all experiments. All experiments were performed in LB Miller broth.

The *bamB* deletion mutant was constructed via backcrossing from the McClelland strain collection (strain C3425, found in ^25^) to SB300 using P22 phage transduction. Resultant kanamycin resistant colonies were sequenced at the *bamB* locus to confirm loss of the coding region. The *bamB* mutant showed no growth defect during *in vitro* growth compared to its wild-type parent (data not shown).

Recombinant secretory IgA (STA121) and its cognate Fab fragment recognizing the O:12-0 O-Antigen serotype was generated as previously described^1,12,26^. IgA and the Fab fragment were fluorescently labeled with AlexaFluor 488 or 568 per the standard protocol of a commercially available antibody labeling kit (Thermo Fisher, A20181 or A20184).

### Atomic Force Microscopy

*Salmonella* Typhimurium from an overnight culture was washed three times in PBS and diluted prior to deposition on polylysine-or (3-aminopropyl)triethoxysilane (APTES)-coated mica. After deposition of a few minutes, the surface was rinsed with PBS to remove loosely bound bacteria. Then after, an aliquot of SIgA was added to the cell layer and allowed to interact for 20 minutes. At this stage, the surface was rinsed with mQ before dying in an air stream. Atomic force microscopy (AFM) imaging was performed using a multimode 8 Bruker AFM in tapping mode using a common silicon AFM tip. Analysis of the images was done with Nanoscope and Gwyddion software. Polylysine-or APTES were purchased from Sigma Aldrich.

### Immuno-labeling of live bacterial cells

100 uL of a *Salmonella* overnight culture was centrifuged and washed 3x with 1 mL PBS. The cell pellet was then resuspended in 100 uL PBS containing fluorescently labeled IgA or Fab. The cells and antibody were co-incubated for 30 minutes at room temperature in the dark. The cells were washed an additional 3x with 1 mL PBS to remove excess antibody and finally resuspended in 100 uL PBS.

### Fluorescence recovery after photobleaching and image analysis

Fluorescently labeled cells were pipetted onto an agar pad and visualized using an Olympus FluoView3000 confocal microscope.

Bleaching of fluorescently labeled antibodies was performed on multiple cells over a circular region of interest (ROI) with a diameter of 11 pixels. The ROI was bleached by setting the laser power to 100% and images of the entire field were collected every 1 s to monitor the kinetics of fluorescence recovery.

Resultant images were analyzed using FIJI (version 2.1.0/1.53c). The Nikon files were converted to a TIFF stack. Fluorescence levels of the ROI were measured over time using the “measure” built-in function. Rate and percent of fluorescence recovery were quantified from the fluorescence curves. Local curvature of the ROI was measured using the built-in “Compute Curvatures” function.

### Live-cell immunofluorescence and image analysis

For experiments investigating stochastic loss of STA121 binding, bacteria bound to fluorescent IgA were loaded into a microfluidic chip using a pipette tip, as described in ^12^. The microfluidic chip was then situated on an Olympus IX81 inverted scope within a incubation chamber heated continuously to 37C. LB Miller medium containing fluorescent IgA-STA121 (1 ug/mL) was flowed through the chip at a rate of 0.2 mL/h using a syringe pump (NE-300, New Era Pump Systems). Phase contrast and fluorescence images were acquired every 3 minutes at 100X.

Antibody wash-out experiments were performed similarly, however after approximately 2 h of growth in Fab or IgA containing (1 ug/mL) medium was exchanged for pure LB Miller.

Image stacks were were deconvolved in MATLAB using a point-spread function ^27^ and subsequently analyzed using SuperSegger ^28^. Images were segmented using *Salmonella* optimized segmentation constants ^29^and manually curated for errors in boundary-calling and tracking.

Per-cell antibody levels were binned as a function of generations and pole-identity from the initial mother cell. Number of cells analyzed per condition are reported in the text. Total fluorescence levels were taken from the mean pixel intensity across an entire cell. Old pole and new pole fluorescence intensities are reported as mean pixel intensity of the old or new pole half, measured from the midpoint of the cell at the last image before division. Consensus images were created by superimposing 5 cells of identical generation-pole-identity with the oldest poles positioned to the left.

### Aggregate breakage experiments

An overnight *Salmonella* wt or *bamB* culture was diluted 1:1000 in 100 uL fresh LB. 4 ug STA121 IgA, Fab, or vehicle control was added to each subculture and incubated for 3 h at 37C (∼ 5 divisions) with no shaking to prompt the formation of enchained aggregates. Cells were centrifuged at low speed to pellet and very gently resuspended in PBS to stall growth and remove excess antibodies. These samples were then either spotted onto an agar pad for imaging or subjected to vortexing at different intensities for 10 s. After vortexing, the cells were spotted onto an agar pad and imaged. For each condition, 10-12 images were taken per experiment, with a total of 3 biological replicates (30-36 fields of view per condition).

Cellular aggregate size was calculated using the native FIJI plug-in. Distributions of aggregate size per strain and per shear force condition were fitted with an exponential decay function and the computed exponential decay constants were compared.

### Statistical and data analyses

Quantitative data were analyzed in R (RStudio, version 1.4.1717).

## Acknowledgements

We are grateful to the members of the Microbial Systems Ecology and Mucosal Immunology groups for all their feedback on this work. We thank Tobias Schwarz and team at ScopeM for help with standardizing the FRAP experiments. This work was supported by ETH SEED to AH and 51NF40_180575, 31003A-16997 to MA. This work was funded by NCCR Microbiomes, a research consortium financed by the Swiss National Science Foundation (ES, MA); Swiss National Science Foundation (40B2-0_180953, 310030_185128) (E.S.), European Research Council Consolidator Grant (865730-SNUGly) (E.S.), Gebert Rüf Microbials (GR073_17) (E.S.); Botnar Research Centre for Child Health Multi-Invesitigator Project 2020 (BRCCH_MIP: Microbiota Engineering for Child Health) (E.S.), and US National Institutes of Health (R01AI165570) (B.M.S.).

---

**Figure S1.**
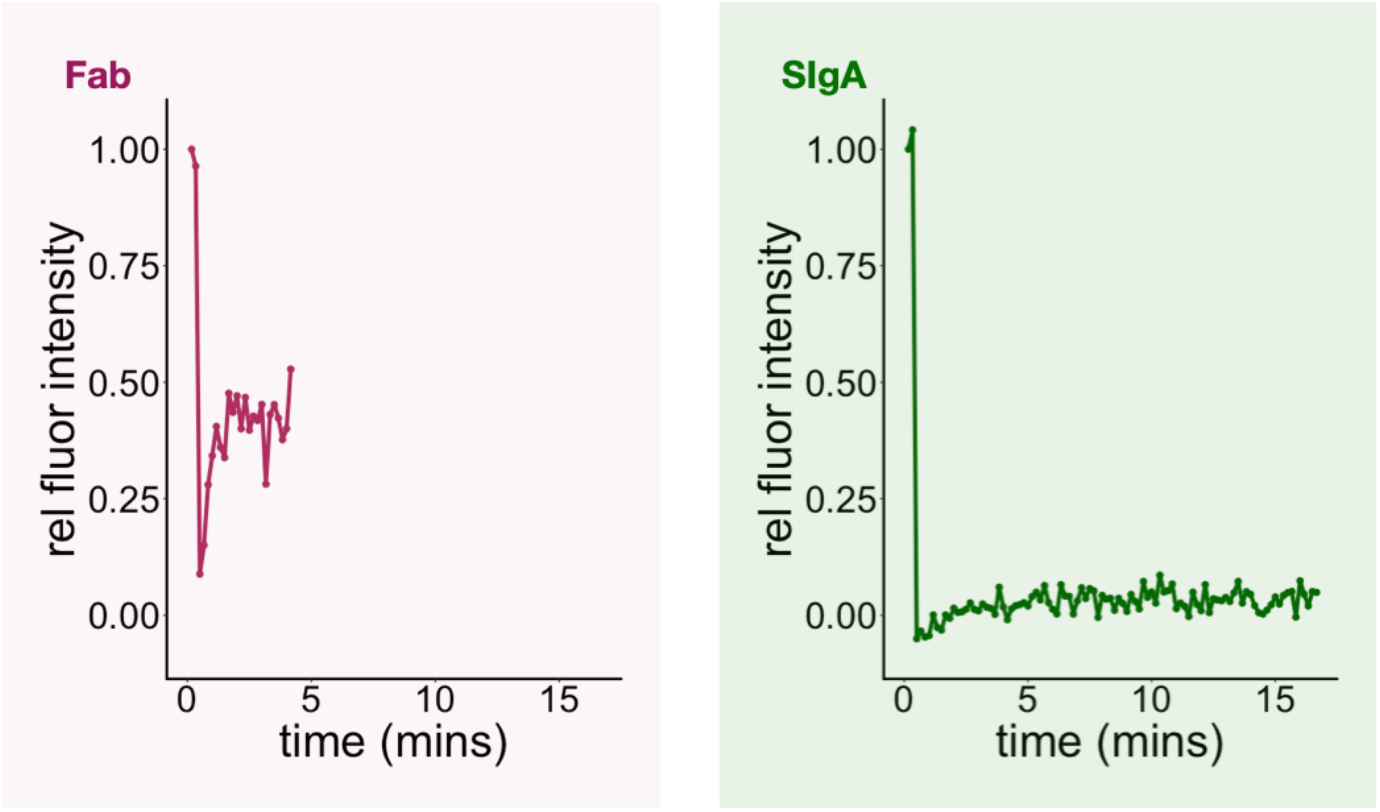
Fluorescence recovery after photobleaching curves. Representative FRAP curves of cells labeled with either the Fab fragment or SIgA.

**Figure S2.**
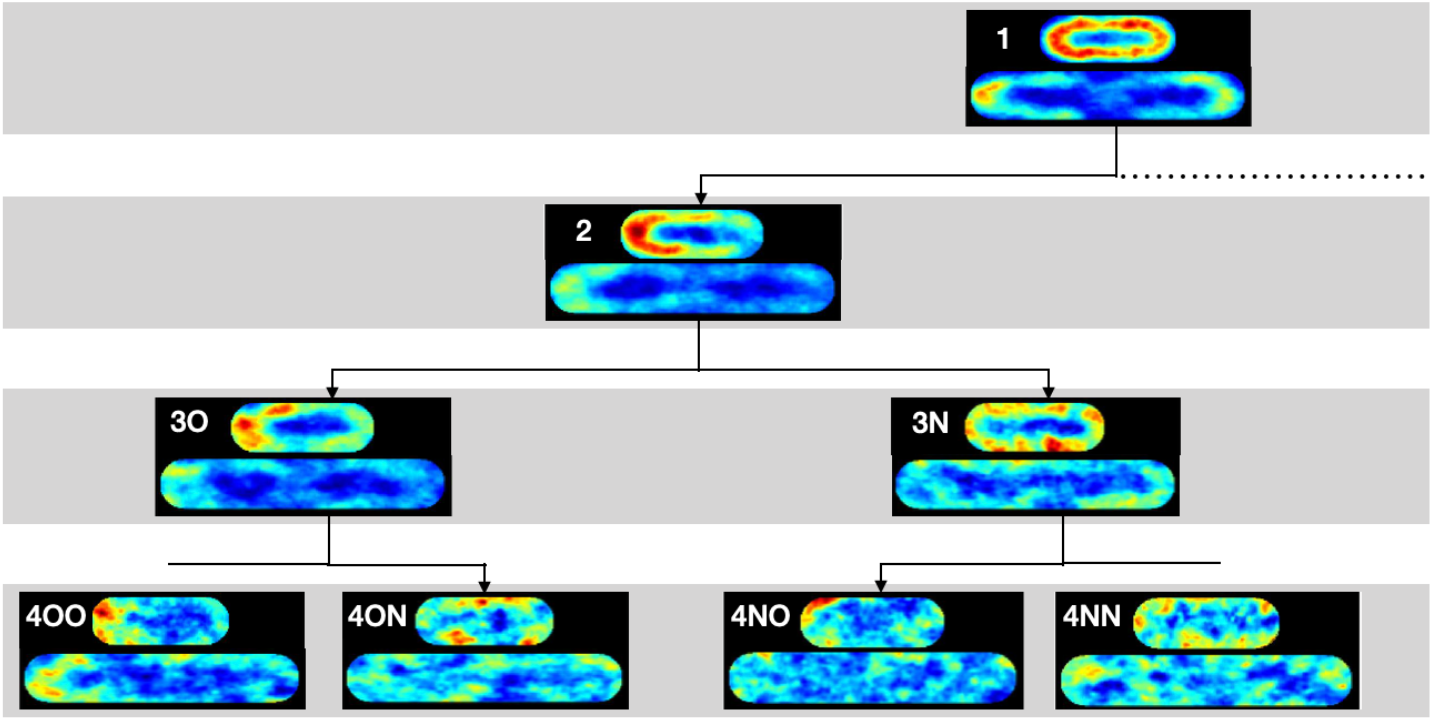
Consensus images of SIgA:O-Ag complexes through *Salmonella* growth. Consensus images of 5 lineages of cells by generation and cellular lineage identity shows distribution of SIgA:O-antigen complexes over time. Top image in each panel shows the cell either as first imaged or the frame directly after division. The lower image in each panel shows the cell one frame before division occurs. Number indicates generation, O for old pole cell, and N for new pole cell. Heatmap levels normalized for each cell identity, not absolute fluorescence levels.

**Figure S3.**
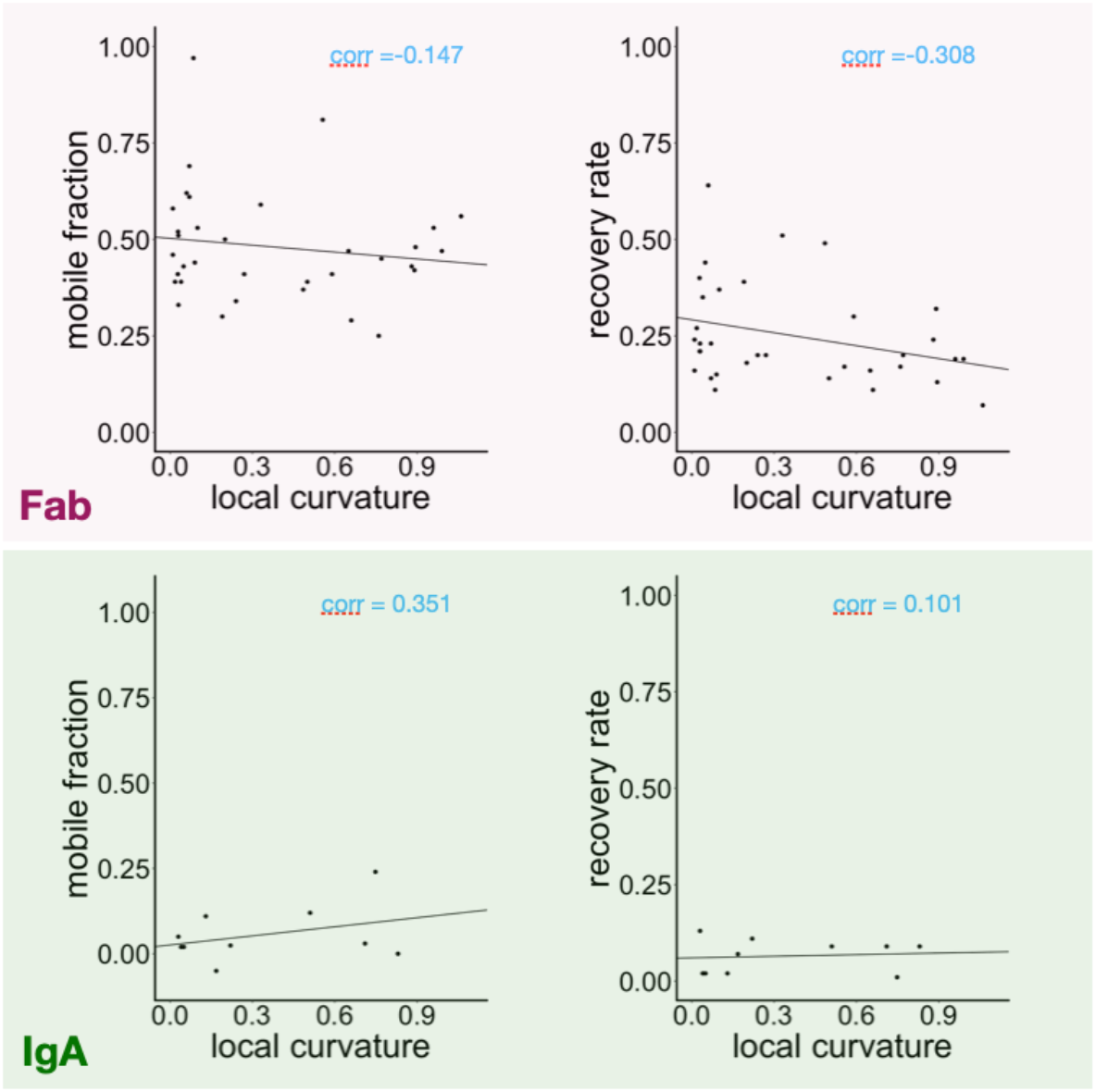
Correlation of local curvature and FRAP parameters. FRAP measurements were taken at different points on the cell. For each photo bleached region of interest, the local curvature was measured and correlated to mobile fraction measured or recovery rate. Corr = Pearson’s correlation

**Figure S4.**
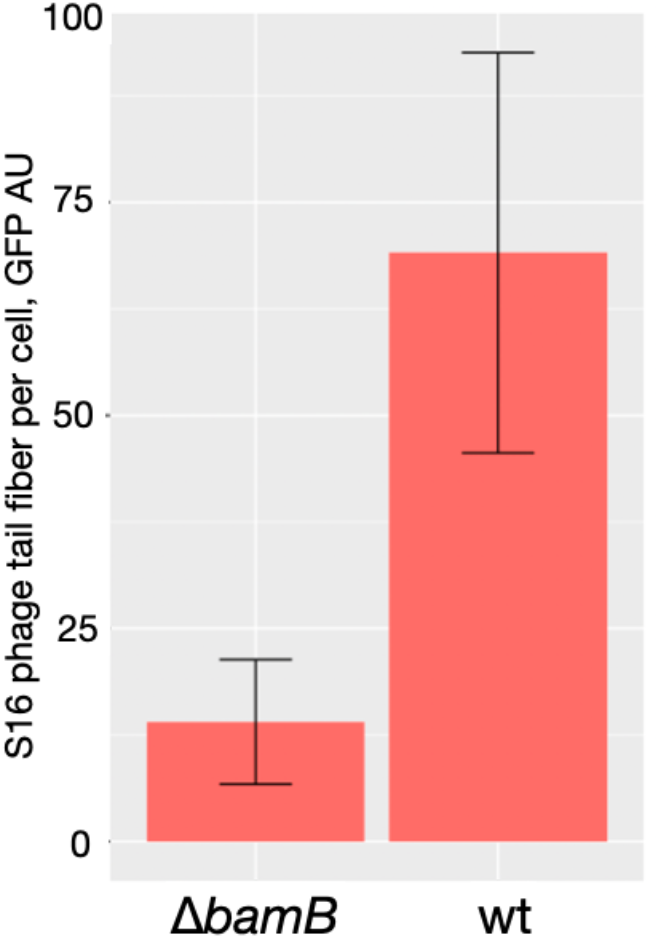
The *bamB* mutant shows lower levels of surface OMPs. Level of surface OmpC in wt and *bamB* cells. Cells were co-incubated with the S16 phage tail fiber specific for OmpC and per-cell fluorescence levels were quantified by microscopy.

**Figure S5.**
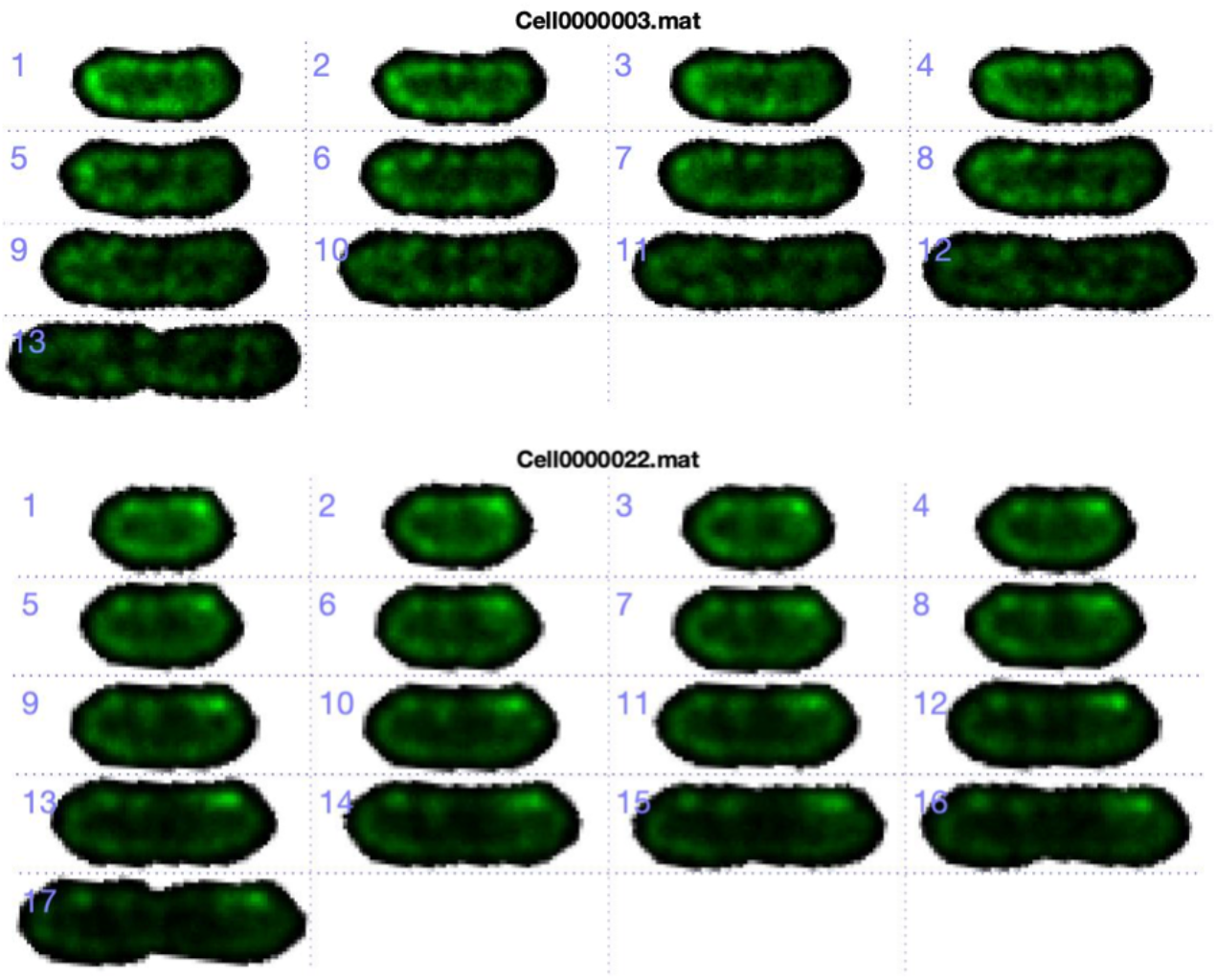
The *bamB* mutant does not show biased inheritance of SIgA. SIgA staining of bamB cells through growth. Green indicates SIgA localization and each number indicates a single cell through growth imaged every 3 minutes.

